# *CalliCog:* an open-source cognitive neuroscience toolkit for freely behaving nonhuman primates

**DOI:** 10.1101/2024.11.26.625450

**Authors:** Jack T. Scott, Bruno L. Mendivez Vasquez, Brian J. Stewart, Dylan D. Panacheril, Darren K. J. Rajit, Angela Y. Fan, James A. Bourne

## Abstract

Nonhuman primates (NHPs) are pivotal for unlocking the complexities of human cognition, yet traditional cognitive studies remain constrained to specialized laboratories. To revolutionize this paradigm, we present *CalliCog*: an open-source, scalable in-cage platform tailored for freely behaving experiments in small primate species such as the common marmoset (*Callithrix jacchus*). *CalliCog* includes modular operant chambers that operate autonomously and integrate seamlessly with home cages, eliminating human intervention. Our results showcase the power of *CalliCog* to train experimentally naïve marmosets in touchscreen-based cognitive tasks. Remarkably, across two independent facilities, marmosets achieved touchscreen proficiency within two weeks and successfully completed tasks probing behavioral flexibility and working memory. Moreover, *CalliCog* enabled precise synchronization of behavioral data with electrocorticography (ECoG) recordings from freely moving animals, opening new frontiers for neurobehavioral research. By making *CalliCog* openly accessible, we aim to democratize cognitive experimentation with small NHPs, narrowing the translational gap between preclinical models and human cognition.

**Motivation:** Cognitive neuroscience research involving nonhuman primates (NHPs) has traditionally been confined to a few highly specialized laboratories equipped with advanced infrastructure, expert knowledge, and specialized resources for housing and testing these animals. The common marmoset (*Callithrix jacchus*), a small NHP species, has gained popularity in cognitive research due to its ability to address some of these challenges. However, behavioral studies in marmosets remain labor-intensive and restricted mainly to experts in the field, making them less accessible to the broader scientific community. To address these barriers, we introduce an open and accessible platform designed for automated cognitive experiments in home cage settings with marmosets. This system supports the integration of cognitive behavioral analysis with wireless neural recordings, is cost-effective, and requires minimal technical expertise to build and operate.

## Introduction

Nonhuman primates (NHPs) are unparalleled among animal models for studying human cognition due to their neurobiological, physiological, and genetic similarities. Recent advances in NHP research, such as in vivo calcium imaging (Bollimunta et al., 2021; Kondo et al., 2018; Wu et al., 2024), large-scale electrophysiology (Liu et al., 2024; Trautmann et al., 2023), and optogenetic techniques (Jendritza et al., 2023; Tremblay et al., 2020), have opened new avenues for investigating the neural circuits underlying cognition. However, despite these technological improvements, there has been relatively little progress in developing reliable and reproducible methods for cognitive behavioral testing in NHPs (Scott & Bourne, 2022). Given the inherent challenges in NHP research, including small sample sizes, high costs, and the need for specialized knowledge, improving the accessibility and reproducibility of cognitive behavioral testing methods is crucial.

Traditionally, cognitive behavior in NHPs is studied by transporting animals from their home cage to a testing environment, where they perform tasks in the presence of a human investigator. These tasks often involve touchscreen interfaces or eye-tracking systems and typically require restraint or physical handling. However, these conditions can introduce stressors that may confound the validity of the data, necessitating time-consuming habituation training. Furthermore, conventional approaches were primarily developed for larger NHPs like macaques, which are more tractable under these conditions. In contrast, smaller NHPs such as common marmosets (*Callithrix jacchus*) and other callitrichids, as natural prey species, tend to be less amenable to restraint and more sensitive to stress, making traditional testing methods less suitable. Likewise, separating callitrichids from their social unit for testing can be more detrimental than for macaques, as they are often group-housed in research facilities due to greater social tolerance and reliance on cooperative breeding.

To address these challenges, the field of NHP research is increasingly shifting toward home cage-based behavioral testing. While home cage testing has been widely used in rodent research, its application to NHPs has gained traction only in recent years (Butler & Kennerley, 2019; Cabrera-Moreno et al., 2022; Calapai et al., 2017; Calapai et al., 2022; Fizet et al., 2017; Ramezanpour et al., 2024; Sacchetti et al., 2021; Tulip et al., 2017; Wither et al., 2020; Womelsdorf et al., 2021). Home cage testing systems allow for more naturalistic behaviors by minimizing external stressors and reducing the time needed for habituation. They can also be fully automated improving experimental throughput and reducing the need for human intervention (Berger et al., 2018; Butler & Kennerley, 2019; Cabrera-Moreno et al., 2022; Calapai et al., 2022; Fizet et al., 2017; Sacchetti et al., 2021; Tulip et al., 2017). Recent studies have also demonstrated the feasibility of integrating wireless electrophysiological implants with touchscreen-based home cage systems, enabling examination of the physiological readouts of cognitive functions in freely behaving animals (Hansmeyer et al., 2023; Wong et al., 2023). Yet despite all these advances, significant barriers remain for the widespread use of home cage-based testing. Current systems often implement different interfaces and test batteries, making it difficult to compare results across studies. Additionally, many home cage testing devices are proprietary or prohibitively expensive and can implement inflexible protocols that are challenging to modify without specialized knowledge. This lack of accessibility discourages the broader use of these methods by scientific community members whose research might otherwise benefit from studying cognition in NHPs.

In this study, we aimed to bridge this methodological gap by developing an open-source platform for studying the neural correlates of cognitive behavior in NHPs. Our system, called *CalliCog* (*<u>Call</u>ithrix jacchus* <u>cog</u>nitive platform), integrates with the housing facilities of marmosets to enable autonomous cognitive behavioral experiments using operant conditioning. Although designed for marmosets, *CalliCog* can be adapted to other small NHP species. Across two facilities with different housing conditions, we demonstrate that *CalliCog* can autonomously train behaviorally naïve animals for touchscreen-based cognitive testing without human intervention. Additionally, we evaluate animal performance in tasks designed to assess behavioral flexibility and working memory and show how *CalliCog* can be integrated with wireless electrocorticography (ECoG) for synchronous neural recording. *CalliCog* is constructed from inexpensive materials and easily fabricated using standard equipment. All resources required to replicate the *CalliCog* system, including hardware inventory, CAD models, source code, and wiring schematics, are freely available.

## Results

### *CalliCog* integrates with primate housing facilities for home cage-based cognitive experiments

*CalliCog* is a flexible and scalable solution for combining cognitive behavioral studies and *in vivo* neurophysiological recordings with cage-housed nonhuman primates. The design of *CalliCog* followed four primary motivations: 1) It should encourage naturalistic, goal-oriented behavior by integrating with the home environment and relying only on voluntary engagement; 2) Behavioral training and testing should be performed autonomously without a human operator, reducing the time allocated to hands-on experimentation; 3) Measured behaviors should be easily synchronized with neural recordings from minimally invasive devices that do not restrict free behavior; and 4) It should embody specific attributes – easily reproducible, low cost, highly flexible and scalable – that enable it to be easily adapted to most housing facilities and laboratories.

*CalliCog* implements a modular design that separates interfaces for acquiring behavioral and neural data from those used to control and monitor experiments. Only two modules are required to perform its minimum functions: an operant chamber that attaches to a home cage and administers experiments; and a computer, typically housed externally to the housing facility, that is used to monitor and control experiments (hereafter referred to as the ‘executive PC’). Each operant chamber is controlled by a local miniature computer within the apparatus (hereafter called the ‘agent PC’), which communicates with the executive PC over a wired network in real time. A unique aspect of *CalliCog* is that the functions of both the executive and agent PCs are intentionally abstracted. While the agent PC’s purpose is to deliver behavioral experiments to an animal and record behavioral responses, the executive PC provides task instructions to each agent PC and receives feedback on behavioral performance. Based on this feedback, the executive PC can autonomously adjust task instructions depending on the user’s experimental design. This bipartite design removes all unnecessary operations from the operant chamber, such as any ‘decision-making’ processes or background programs, that would otherwise increase the computational load on the agent PCs and impact their performance. Notably, a single executive PC may control many agents depending on the scale of a given housing facility, allowing for high-throughput testing of multiple cage-housed animals simultaneously.

Here, we demonstrate the operations of *CalliCog* for unsupervised training and testing of several marmosets housed in either single- or pair-housed home cages using multiple operant chambers **(Figure 1).** During experimentation, chambers are affixed to the front of cages using custom-built brackets, allowing an animal to enter the chamber at any time voluntarily. Animals engage with behavioral tasks by interacting with stimuli presented on a capacitive touchscreen (779GL-70NP/C/T; Lilliput Electronics), and liquid reward (dissolved marshmallow in water, 20% w/v) is dispensed through a reward spout for positive reinforcement using a reward module driven by an Arduino microcontroller (Arduino Uno Rev3; Arduino). Live video is streamed wirelessly using an integrated Raspberry Pi camera module (Raspberry Pi Zero W with Raspberry Pi Camera Module 2; Raspberry Pi Foundation) to the executive PC (M2 Mac Mini; Apple Inc.) for uninterrupted surveillance. The agent PC (NUC 13; Intel Corporation) in each operant chamber is directly connected to an isolated local area network (LAN) via a network switch (GS116NA; Netgear, Inc.) and wired router (RV160; Cisco Systems, Inc.) placed centrally in the housing facility. In our facility, we contained the switch and router in a ‘mobile deployment hub’, a custom-built cart that was used to transport operant chambers into the area and provide them with inputs to wired ethernet and power **(Figure S1A)**. The network switch in the mobile deployment hub is connected directly to the executive PC housed in a laboratory adjacent to the housing facility. Notably, while the mobile deployment hub streamlined operations in our housing facility, it is not essential. Only a typical mains power source and an ethernet link between PCs are required for *CalliCog* to function.

**Figure 1.**
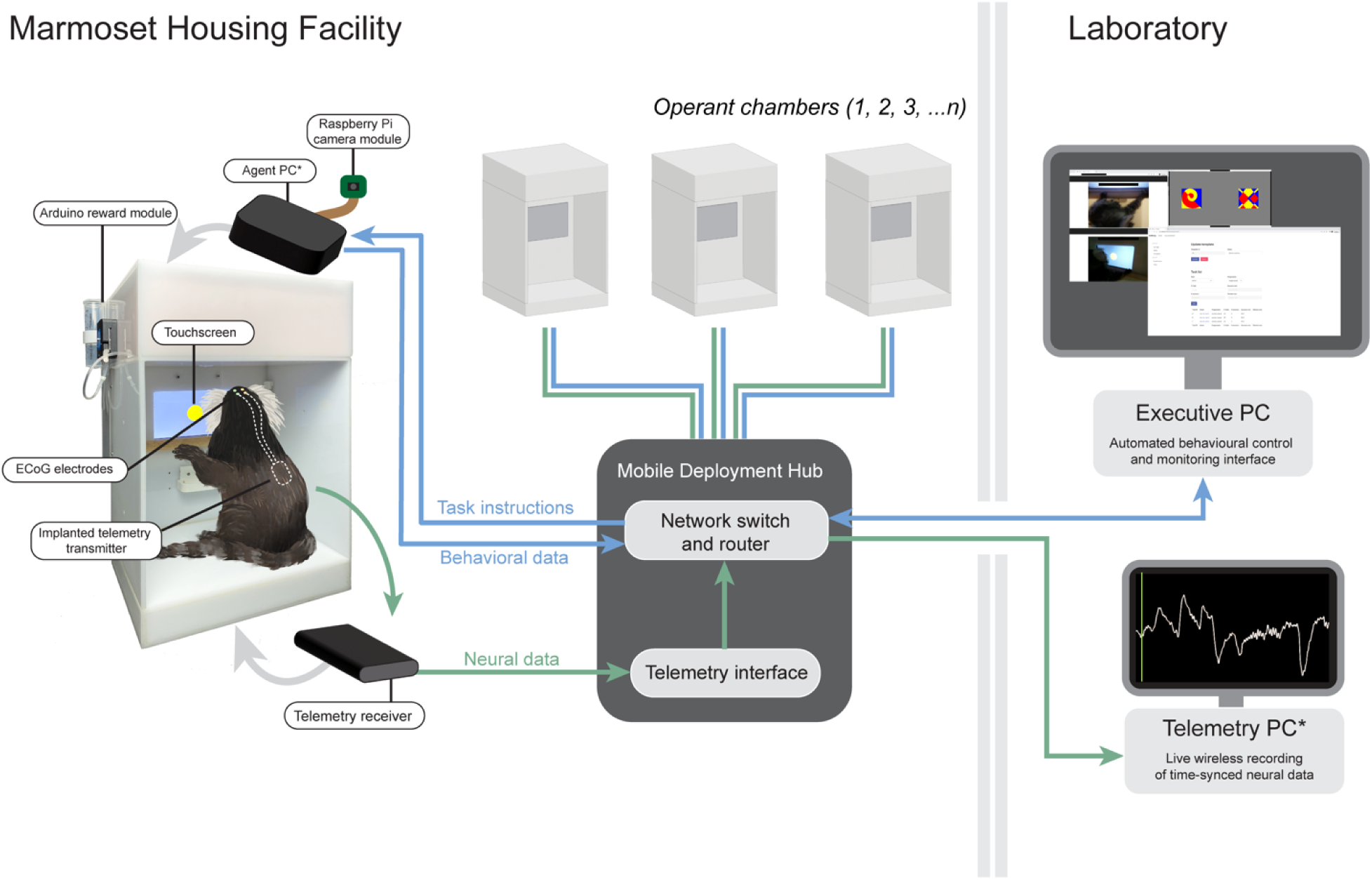
High-throughput, automated cognitive experiments in a marmoset housing facility using *CalliCog.* Individual home cages are fitted with operant chambers (inset image), which are used to train and test behaviorally naïve animals in touchscreen-based cognitive tasks using positive reinforcement. The functions of each operant chamber are controlled by an internal agent PC, which receives trial instructions over a local network from an executive PC housed outside of the housing area. The executive PC receives behavioral data from the agent PC and queries performance against defined parameters to determine automated task progression. A single executive PC can simultaneously administer multiple operant chambers in a housing facility, facilitating a flexible and scalable approach to home cage testing. In addition, devices for neural recording can be integrated with the local network to study the neural correlates of cognitive behavior. In this example, an off-the-shelf system for wireless telemetry recording (indicated by green arrows) is used to acquire ECoG via wireless receivers while animals engage in touchscreen-based tasks. **\***Agent PCs are synchronized to a clock set by the telemetry PC using a conventional network time protocol, allowing for synchronous behavioral and neural data timing.

*CalliCog* is compatible with techniques used to study behavior-related neural activity *in vivo* by synchronizing behavioral timestamps with neural recordings. Here, we demonstrate the feasibility of such an approach by integrating an ‘off-the-shelf’ radiotelemetry system into the *CalliCog* network to record wireless ECoG while marmosets engage in behavioral tasks (**Figure 1**). Using subcutaneous telemetry transmitter implants (HD-S02, Data Sciences International) that record from intracranial electrodes, neural activity can be wirelessly transmitted from an implanted animal to a telemetry receiver (RSC-1; Data Sciences International) located in a compartment beneath the operant chamber. This signal is then transmitted through a telemetry interface (MX2; Data Sciences International) to an external computer for recording (ThinkPad P12s Gen 1; Lenovo) (hereafter referred to as the ‘telemetry PC’). To ensure that neural recordings can be precisely aligned with behavioral events, we configured the telemetry PC to act as a network time protocol (NTP) server to synchronize timings with each agent PC. By intermittently querying the offset between PC clocks via the NTP daemon on the agent PC during routine use, timings were determined to be reproducibly synchronized within 12-15 ms. As a result, neural activity and behavioral responses could be accurately aligned in later offline analysis with consideration to this error margin. Importantly, while we demonstrate the utility of *CalliCog* here with ECoG telemetry, this principle of network timing could feasibly be generalized to a range of devices for neural recording or stimulation.

### Operant chambers are designed for easy construction and use minimal, low-cost resources

The *CalliCog* operant chamber is a bespoke testing interface designed specifically for cognitive behavioral experiments in marmosets. To be rapidly and tractably adopted by other laboratories, it uses primarily inexpensive and readily available components (a complete inventory of materials and their specifications can be found via the DOI in the key resources table). Moreover, building the foundations requires only flat panels of opaque acrylic or polyvinyl chloride (PVC) held together by an epoxy adhesive, and a minimal number of fastenings and fixings to attach removeable lids. The remaining inserts can be readily made using standard 3D printers found in many laboratories. All computer-aided design (CAD) files for fabrication and 3D printing are provided online (see key resources table for DOI).

A total of 4 operant chambers were used concurrently in the current study. Each operant chamber contains four waterproofed compartments: the internal testing chamber, an upper housing for the agent PC and electronic components, a rear housing for the touchscreen, and a lower housing for placing a telemetry receiver. The internal testing chamber houses two 8-megapixel Raspberry Pi cameras for video surveillance, one in the center of the ceiling for a top-down view of the chamber, and another (optionally) housed in a 3D-printed insert at the front. The rear of the chamber houses a 7” capacitive touchscreen supported by 3D-printed brackets, a reward spout, and a custom 3D-printed tray for delivering liquid reward. In addition, 3D-printed braces support a height-adjustable, wooden ‘training bar’ for animals to grasp while they perform behavioral tasks **(Figure S1B)**. While the training bar was usually positioned below the touchscreen, it could be used to direct animal responses during training depending on user preference. For example, during early training, if animals touched the display by accident, training bars may be used to block unintentional touches and ensure that responses are volitionary. In routine training and testing, implementing the training bar below the touchscreen at all other times encouraged engagement with the touchscreen for extended periods. This was likely because unsupported rearing on hindlimbs is an unnatural posture for marmosets, as arboreal animals, to maintain over time.

At the top of the operant chamber, the electronics housing contains the agent PC that controls peripheral electronics **(Figure S1C)**. While we mainly used the Intel NUC 13 model in this study, system requirements can theoretically support any miniature PC capable of running the Ubuntu operating system (version 22.04) (for devices with confirmed functionality, see the inventory in the repository listed in the key resources table). The agent PC controls the touchscreen display and records screen touch responses. In addition, trial outcomes in behavioral tasks (i.e., correct or incorrect responses) trigger the agent PC to communicate with a reward module, which controls the emission of auditory cues and the dispensation of liquid reward. The reward module consists mainly of an Arduino microcontroller with a custom shield, an audio speaker (C0604B; Altronic Distributors Pty Ltd.), a stepper motor (ROB-09238; Sparkfun), and a driver (Big Easy Driver; Sparkfun). The stepper motor and a 60 ml syringe, which provides a reservoir for liquid reward, are mounted externally on the side of the operant chamber. The wiring schematic and list of components for the reward module is available in the online repository (see key resources table for DOI).

Critically, the operant chamber’s upper and rear compartments are lined with copper foil tape for electrical shielding, which is anchored to a common ground with the electronic components. This approach successfully mitigated electromagnetic interference from the electronics within the testing chamber—particularly the touchscreen—which otherwise interfered with wireless transmission from radiotelemetry implants and interrupted signals during recording.

### Open-source software allows for both automated training protocols and flexible manual experimentation

The *CalliCog* software is primarily based on a combination of custom scripts and third-party, open-source Python packages. Most notably, core processes related to the presentation of stimuli and detection of responses in the agent PC are generated by PsychoPy, a powerful open-source toolbox for running psychophysics experiments (https://psychopy.org/) (Peirce et al., 2019). Overarchingly, the *CalliCog* software is functionally distributed between the agent PC and executive PC, with the former controlling all local operant chamber operations and the latter handling experimental workflows, data storage, querying, and live monitoring. Ease of use, flexibility, and accessibility were primary considerations in the design of the architecture, such that even a user with minimal programming knowledge could design and run custom-built behavior experiments. In addition, experiments can either be performed with manual control or using automated protocols, depending on user requirements. Below, we summarize the software’s main features and describe how protocols are designed and executed. However, a more comprehensive description, including a guide for installing and creating custom experiments, is available along with the source code in a public GitHub repository (https://github.com/NIMH-SCCN/callicog).

The executive PC provides the interface for a user to design, conduct, and monitor behavioral experiments. Executive PC installation is currently optimized for the Mac operating system, and in our setup, the executive PC was an Apple M2 Mac Mini running macOS Sonoma (version 14.5). The user can initiate and control an experiment on a given operant chamber on the executive PC by opening a new terminal window and entering simple commands built into the *CalliCog* software (refer to GitHub repository for instructions). In addition, the executive PC can be used to monitor active experiments. Each operant chamber acts as a virtual network computing (VNC) server for screen sharing on boot, allowing the user to visualize and interact with a real-time mirror of the display presented on the touchscreen. Video streams from operant chambers are broadcast wirelessly from individual Raspberry Pi camera modules using a freely available third-party tool, RPi Cam Web Interface (version 6.6.15) and monitored through a web browser on the executive PC. Importantly, video access is not limited to the executive PC and can be password-enabled and streamed over any wireless network of choice (e.g., the public network of an academic or research institute). This allows for remote monitoring of automated experiments from outside the housing area using devices such as laptops or smartphones.

The executive PC also houses a custom web application (web app) that can be used to monitor behavioral data and generate reports for offline analysis. The web app acts as an interface for a relational database, implemented in PostgreSQL, that stores all experimental data and parameters. The database is structured as a series of individually timestamped events from an experiment. These events describe all parameters related to what is presented to a test subject (e.g. stimulus attributes, display timings) and the subject’s responses (e.g. screen touch coordinates, the timing of screen touch and release, response latencies, performance outcome). Each event is associated with a unique identifier that can be traced to the exact screen window on which it occurred in a specified trial, within a given session of trials, during a particular experiment. Based on this database organization, reporting functions in the web app can be implemented to navigate and query large experimental datasets with ease.

The design of experimental protocols to be run by the executive PC is also performed within the web app environment, and for its basic usage does not require prerequisite programming knowledge. Protocols are written by the user into the database using the web app, and these protocols are structured hierarchically to facilitate flexibility in experimental design (for complete details and instructions on designing experiments using *CalliCog*, refer to the online GitHub repository). By navigating this hierarchical structure, a user may configure experimental parameters at multiple levels, from the specific attributes of touchscreen stimuli and the design of individual trials to the sequence of individual task stages within a greater protocol to be executed over more extended periods (e.g., days or weeks). When designing a protocol, the user defines an experiment as a sequence of tasks for an animal to be progressed through sequentially (for example, the several stages of a touchscreen training protocol). These tasks are associated with progression criteria that determine the level of performance an animal must achieve to progress to the next task. The capacity of *CalliCog* to evaluate progression criteria based on behavioral performance to progress animals through tasks forms the basis of its capacity to run automated experiments. There are 3 types of progression criteria currently implemented in *CalliCog*, as follows:

- ‘Target-based’: Progression occurs after a set number of trials are performed, regardless of the trial outcome (success or fail). Example: “50 trials”.
- ‘Session-based’: Progression occurs after a consecutive number of sessions (i.e. groups of trials) are performed to a specified level of proficiency. Example: “> 80% success in 3 consecutive sessions of 50 trials”.
- ‘Rolling average’: Progression occurs after a specific proficiency has been achieved over a rolling window of multiple trials, continuing in perpetuity until the criterion is reached. Example: “90% success within the last 100 trials”.

Each of the progression criteria provides different utility depending on the experimental design. For example, ‘target-based’ is implemented when the user desires manual (non-automated) training. Conversely, ‘session-based’ is helpful for contexts where animals should automatically progress to subsequent tasks upon reaching stable levels of performance (e.g., training protocols), and ‘rolling average’ is advantageous for adapting experiments to behaviors that can change at the level of an individual trial (e.g., in learning or decision-making paradigms). To implement these criteria in real-time, the executive PC receives feedback from agent PCs after each trial is executed and compares the current state of performance to specified progression parameters **(Figure 2).** On a trial-by-trial basis, data is sent from an agent PC to the executive and logged in the database. The executive PC then queries the database to evaluate past trials against the progression criterion. Once the executive PC evaluates the most recent performance data against the progression criterion, it sends instructions for a new trial from either the current or subsequent task to the operant chamber. Packaged in JSON text format, the agent PC parses these trial instructions and executes the ensuing trial. These processes occur almost instantly during standard operation and may be performed concurrently across multiple operant chambers and animals.

**Figure 2.**
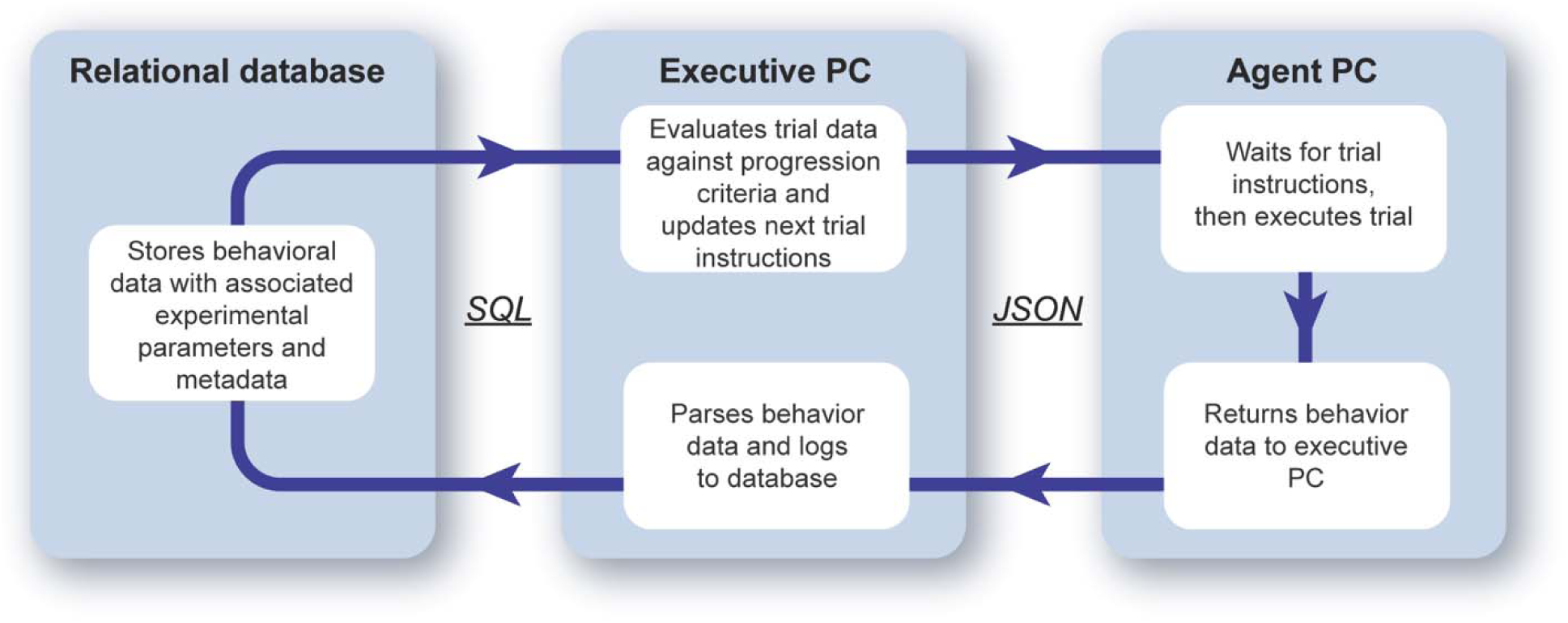
Workflow for automated progression during behavioral training or testing. Once a protocol has been designed and initiated by the user at the executive PC, behavioral experiments run autonomously by evaluating animal performance on a trial-by-trial basis. The executive PC sends trial instructions to an agent PC using the JSON text format, which the agent PC parses for trial execution. Trial data, including all timestamps of display events and touch responses, are then returned to the executive PC and logged via the SQL programming language to a local relational database (PostgreSQL). The executive PC then evaluates the logged trial data against predefined ‘progression criteria’ to determine whether to continue the current task or progress to the next in sequence. Responding to animal performance at single-trial resolution allows the system to adapt tasks based on cognitive behaviors that can change on a single moment (e.g. learning or decision-making).

### Cognitive training and testing of behaviorally naïve marmosets using an automated protocol

To demonstrate the utility of *CalliCog* for autonomous training of cage-housed primates, we introduced a cohort of marmosets (*n* = 8; 3 female, 5 male) to a pipeline of protocols designed for touchscreen training and cognitive testing. All animals selected were in adulthood (age during training: 12 – 72 months) and were naïve to behavioral experiments. The cohort was split between two individual housing facilities with different husbandry procedures and cage housing designs, with 4 animals (M2, M3, M7, and M8) housed in ‘Facility 1’ and 4 animals (M1, M4, M5, M6) housed in ‘Facility 2’ (see STAR Methods for further information). Not all animals in the cohort were subjected to every experiment in this study, as several were taken offline for use in other projects (for details, see **Table S1**). However, notably no animals were removed from the study due to an inability to operate the touchscreen or perform tasks.

Marmosets were progressed through an automated pipeline consisting of individual training or testing protocols with multiple phases **(Figure 3A).** Each protocol contained specific progression criteria defining when animals should progress through constituent phases and subsequent protocols. To summarize the pipeline, animals were first presented with a touchscreen training protocol to learn to interact with visual stimuli for liquid reward. Once proficient in responding to stimuli of variable color and position, animals performed discrimination training to distinguish between correct and incorrect stimuli using feature-based learning. Subsequently, animals performed novel discrimination and reversal learning tasks designed to test the capacity for behavioral flexibility. Finally, they were trained to perform a feature-based working memory task (delayed match-to-sample task (DMTS)), which was then used to test working memory. Discrimination training was intentionally performed before the DMTS, as previous work has shown that training marmosets to attend to stimulus features improves learning outcomes (Nakamura et al., 2018).

**Figure 3.**
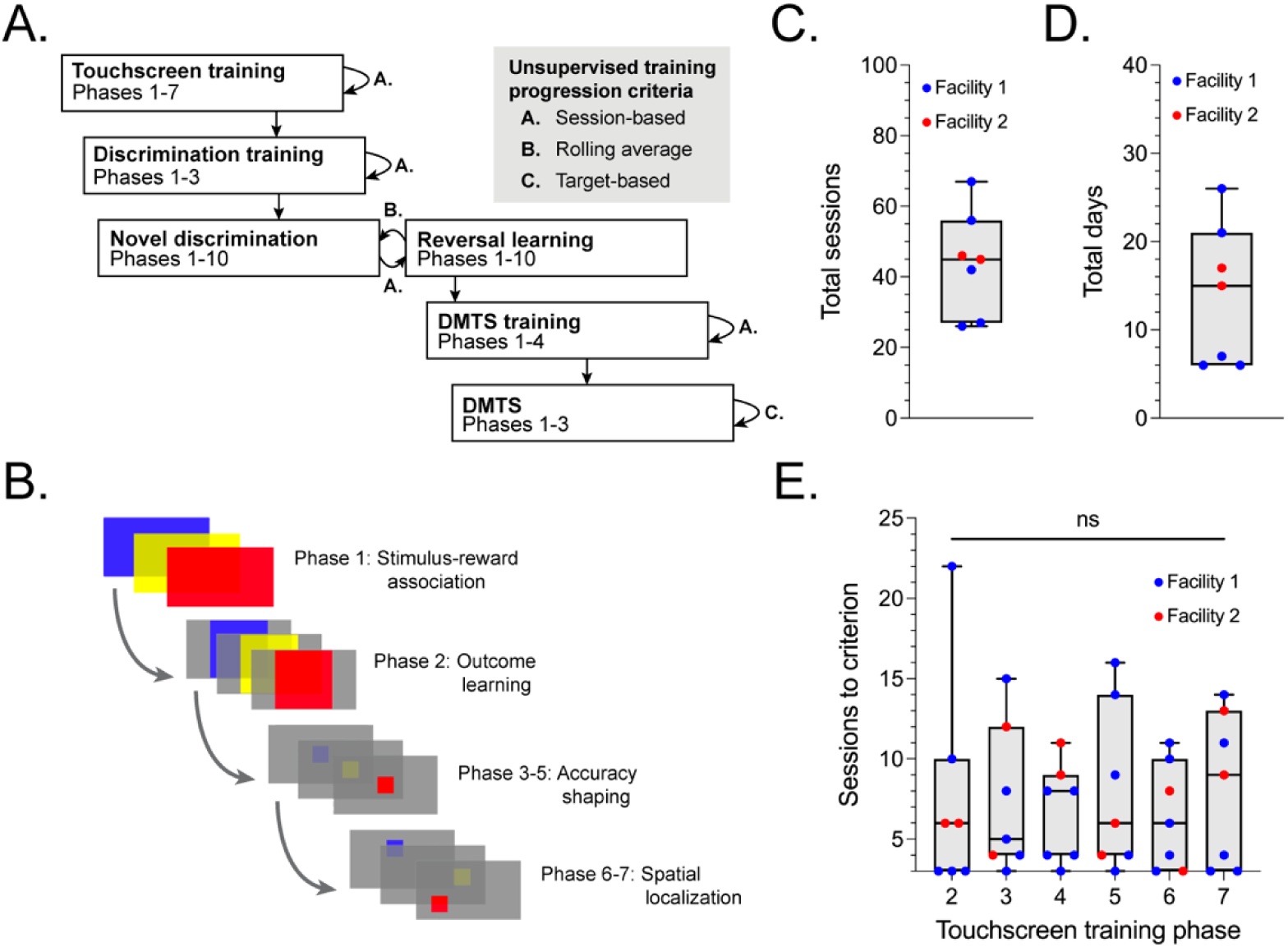
Experimental pipeline and animal performance in automated touchscreen training. **A.** Summary of the unsupervised experimental pipeline, including all stages of training and testing, used in the current study. Each experiment incorporated multiple phases through which animals progressed after achieving predefined progression criteria. Three types of progression criteria were used: ‘session-based’, where animals performed consecutive sessions of trials to a predefined rate of success; ‘rolling average’, where animals performed to a predefined success rate over a rolling window of trials; and ‘target-based’, where animals completed absolute numbers of trials before progression, regardless of successful or failed outcome. **B.** The touchscreen training protocol included 7 individual phases in which animals were gradually trained to respond to small (250 x 250 px) stimuli regardless of color and spatial position. **C, D.** Summarized performance of the experimental cohort over all 7 phases of touchscreen training. The data is presented to show the variability between animals that were housed in different animal facilities. On average, animals completed touchscreen training in 44.1 sessions or 14 days. **E.** Variability between the performance of individual animals on phases 2-7 of the touchscreen training protocol. Performance across individual phases did not vary significantly (*p* = 0.97; Kruskal-Wallis test). Note that animals require a minimum number of 3 sessions to complete each phase.

The touchscreen training protocol was primarily adapted from previous studies (Nakamura et al., 2018; Takemoto et al., 2015) and consisted of 7 total phases **(Fig 3B)**. In phase 1, animals learn to associate screen touches with reward by interacting with a large stimulus, pseudorandomly sampled on each trial as one of three colors (yellow, blue, or red). In phase 2, animals learn trial outcomes by selecting a stimulus contingent with a reward or a grey background contingent with a penalty timeout period (2 s). In phases 3-4, the accuracy of touchscreen responses is shaped by gradually reducing the stimulus size. Finally, in phases 6-7, stimuli appear in progressively greater radial coordinates from the center of the screen so that animals respond regardless of spatial position. Across the entire cohort, all animals completed phases 1-7 over a mean of 44.1 sessions (*SD* = 14.7) and 14 days (*SD* = 8.0) (**Figure 3C and 3D).** The data was similarly distributed regardless of whether animals came from Facility 1 or Facility 2, and no individual animal was considered an outlier in the sample distribution (Z-scores within the range -2 to 2). Between individual phases 2-7 (i.e., once animals had learned to associate screen touches with reward), there was no clear trend in sessions to criterion between animals housed in either facility, and several animals completed phases in the minimum 3 sessions required to reach criterion **(Figure 3E).** There was no main effect of individual training phases on the performance of the animals (Kruskal-Wallis test; *H*(5) = 0.84, *p* = 0.974), suggesting that no individual phase in the sequence was significantly more difficult to perform.

Upon completion of touchscreen training, animals were trained to perform the novel discrimination and reversal learning tasks **(Figure 4A).** In these tasks, animals touch a cue stimulus to initiate a trial. Then, they must select between 2 novel geometric stimuli: one contingent with a reward and the other a penalty timeout. During training, animals perform this task with 3 pairs of novel stimuli and must achieve the progression criterion for each before progressing to the next pair. During testing, 10 stimulus pairs were used for these novel discriminations, each with an associated reversal learning stage. This stage occurs immediately following an animal achieving progression criterion during novel discrimination and involves a reversal of the stimulus-reward contingencies (i.e. the correct stimulus is now the incorrect stimulus, and vice versa). Animals must learn the new contingencies to criterion before progressing to the next stimulus pair. Analysis of the proportions of trials required to meet criterion during reversal learning provides quantitative measures of behavioral flexibility. To evaluate the success of discrimination training, we first examined the number of sessions animals required to reach criterion for each stimulus pair during the discrimination stage. As anticipated, performance improved and became less variable over the 3 stimulus pairs in the training stage as the animals learned the task **(Figure 4B)**. Next, for the 10 stimulus pairs used in the subsequent testing stage, we then compared discrimination performance between the 10 stimulus pairs used for testing to ensure that performance stable and that no stimuli were outliers (i.e. more or less difficult to discriminate than others) **(Figure 4C)**. Contrarily, we detected a main effect of stimulus pair on sessions to criterion (Kruskal-Wallis test; *H*(9) = 17.86, *p =* 0.037) and a greater number of sessions required for stimulus pair 9 compared to stimulus pair 3 (Dunn’s test; *p* = 0.015). Based on this, we determined stimulus pair 9 to be a likely outlier for influencing animal performance, which was further supported by computing the group-wise Z-score for stimulus pair 9 (Z-score = 2.47). As these stimuli were likely to confound reversal learning, we omitted stimulus pair 9 from all subsequent analyses. For reference, images of all stimulus pairs used in these experiments can be found in the code repository listed in the key resources table. Stimulus pair 9 consisted of two images that both contained radial geometry, which likely contributed to discrimination difficulties.

**Figure 4.**
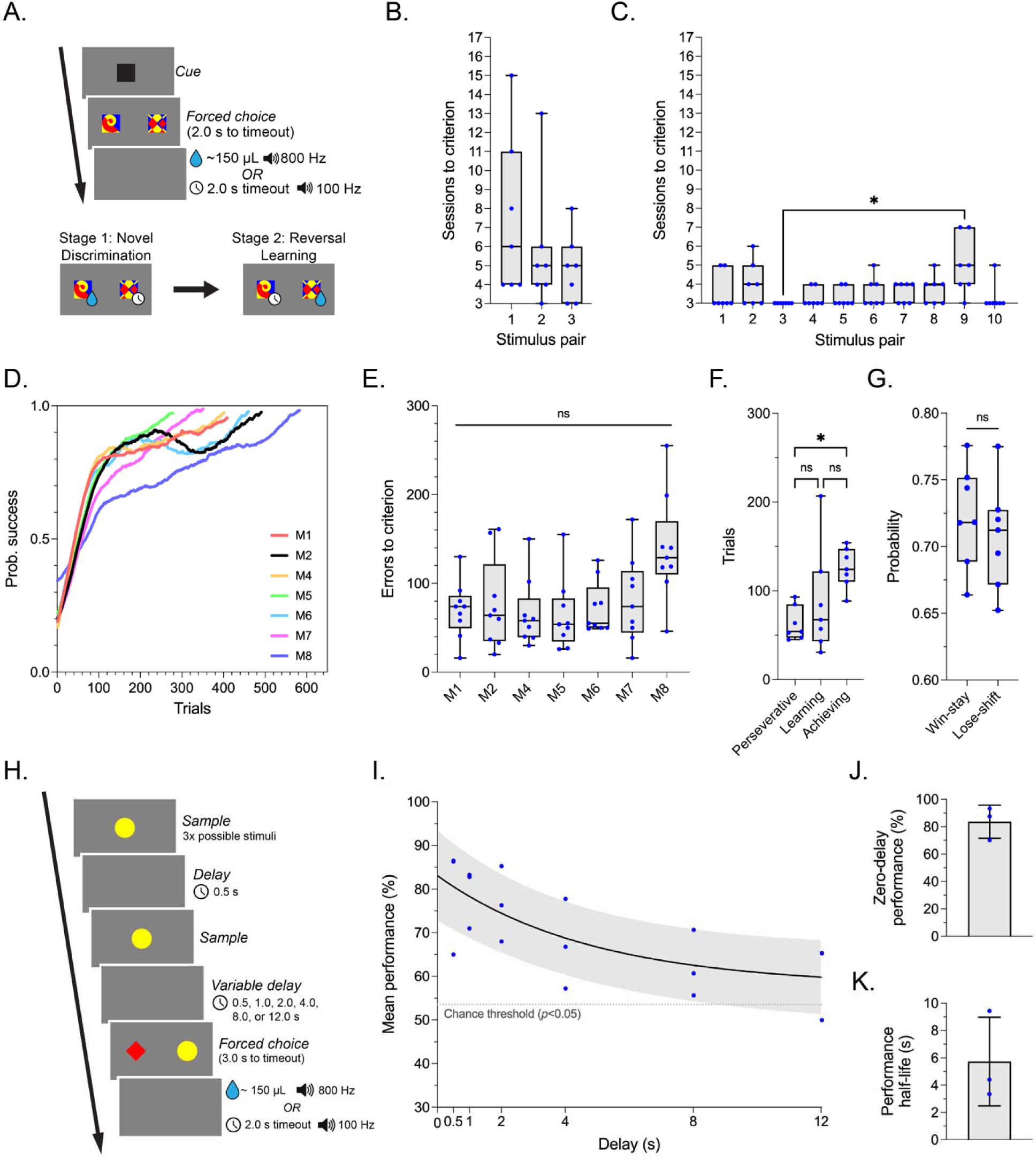
Animal performance in behavioral flexibility and working memory tests using *CalliCog.* **A.** Trial representation of the novel discrimination and reversal learning tasks. Animals interact with a cue to initiate a trial. Then, they must discriminate a rewarded stimulus from a non-rewarded stimulus from a pair of novel stimuli, pseudorandomly presented in left and right positions. Per stimulus pair, animals perform a novel discrimination stage to proficiently learn the stimulus-reward contingencies, which are then instantly reversed in the reversal learning stage after a progression criterion is reached. Behavioral flexibility is evaluated based on performance in adapting responses during reversal learning. **B.** Performance on the 3 stimulus pairs used for novel discrimination training. **C.** Performance on the 10 stimulus pairs used for novel discrimination testing. Stimulus pair 9 significantly contributed to a main effect of stimulus pair on performance (Kruskal-Wallis test with Dunn’s test, *p* = 0.015), so was omitted from subsequent analysis. **D.** Averaged learning curves of individual animals across all 10 reversal learning experiments. The data is smoothed for visualization purposes using a 2^nd^ order polynomial fit to a rolling window of 100 neighbors. **E.** Individual performance in reversal learning experiments between animals, as characterized by the number of errors committed before reaching criterion. The mean performance did not differ significantly between animals (*p* = 0.087; Kruskal-Wallis test). **F.** The proportion of trials animals performed across 3 phases of reversal learning, as calculated from learning curves of cumulative performance using a method for learning curve analysis. Animals performed significantly more trials in the achieving phase than in the perseverative phase (Kruskal-Wallis test with Dunn’s test; *p* = 0.015). The learning phase, in which animals adapted their responses to the new contingencies more consistently, was the most variable. Data points represent the mean number of trials per animal across all 9 reversal experiments. **G.** The probability of animals using win-stay or lose-shift strategies during the learning phase of reversal learning represents the likelihood of correctly adapting responses based on positive and negative feedback, respectively. No significant differences were observed between either strategy (Mann-Whitney U test; *p* = 0.535). **H.** Trial representation of the delayed match-to-sample task (DMTS) used to assess working memory maintenance. Animals are presented with a sample stimulus to be recalled, pseudorandomly selected from a yellow circle, red diamond, or blue star. After interacting with the sample twice, a variable delay period of between 0.5 – 12 s is presented, and animals must then choose the sample in the presence of a distractor to perform the task successfully. The relationship between successful performance and the length of the delay period indicates the capacity for maintaining working memory over time. **I.** The average working memory decay curve generated from the averaged performance data of animals (*n* = 3) performing the DMTS. The data is fit to a simple exponential decay function, and the chance threshold is the performance value at which the percentage of success is significantly greater than the chance performance of 50%. **J, K.** Derived metrics of zero-delay performance and performance half-life from the working memory decay curves of individual animals, which represent the baseline ability to perform the task and the rate of working memory decay, respectively. * *p* < 0.05

Next, we examined individual differences between animals during reversal learning. First, we compared individual learning curves calculated from the average success of each animal per trial across all 9 considered experiments **(Figure 4D).** Learning was largely congruent as the trial number increased, with most animals 80% likely to respond successfully by 200 trials and completing all reversal experiments within 500 trials. The exception to this was animal M8, which reached stable performance levels at a comparatively greater trial number. Next, we evaluated the capacity of animals for behavioral flexibility by quantifying the number of errors committed before reaching criterion. Across the cohort, animals achieved criterion after committing a grand mean of 81.3 errors (*SD* = 26.0). Between individual animals, there were no significant differences in mean errors to criterion between animals (Kruskal-Wallis test; *H*(6) = 11.03, *p* = 0.087), despite animal M8 trending towards greater numbers of errors **(Figure 4E)**. After establishing the base performance in reversal learning, we characterized learning further by generating learning curves for all reversal learning data. A learning curve analysis (see STAR Methods) was then applied to quantify the number of trials that animals performed over 3 phases of learning: ‘perseverative’ (the animal is responding to the previous contingencies), ‘learning’ (the animal is adapting its responses), and ‘achieving’ (the animal has learned the new contingencies). On average, animals performed significantly more trials during the achieving phase compared to the perseverative phase (Kruskal-Wallis test; *H*(2) = 8.51, *p* = 0.009; Dunn’s test; *p* = 0.015) (**Figure 4F**). No significant differences were observed between the number of trials performed in the learning compared to the other 2 phases (Dunn’s test; *p* > 0.05 for both comparisons). To examine the strategies that animals employed during learning, we then calculated the likelihood of successful trials immediately following a success (‘win-stay’) and successful trials immediately following a failure (‘lose-shift’) during the learning phase. Overall, there was no difference between the percentage of either win-stay or lose-shift strategies during the learning phase (Mann-Whitney U test; *U* = 19, *p* = 0.535), indicating that marmosets used both positive and negative feedback equally to respond successfully during learning **(Figure 4G).**

After completing reversal learning experiments, animals progressed to training for the DMTS, a task used to assess working memory **(Figure 4H)**. On a given trial of the DMTS, animals first interact with a ‘sample’ stimulus to be held in memory, pseudorandomly sampled to be one of 3 possible stimuli (yellow circle, red diamond, or blue star). After touching the sample twice, animals are presented with a pseudorandomly sampled delay period (0.5, 1, 2, 4, 8, or 12 s). Immediately following the delay, the animal is presented with the sample stimulus and a distractor selected from the remaining 2 stimuli, and the sample must be chosen correctly to receive a reward. The capacity for maintaining working memory is then assessed by examining successful performance across the delay conditions as a function of time. Due to allocating several animals in the cohort to other projects after reversal learning, only a subset (*n* = 3; M1, M2, and M3) underwent DMTS training and testing. During training, animals are progressed through 4 individual stages in which a simpler version of the task is made progressively more complex (see STAR Methods). During testing, only shorter delays (0.5 – 4 s) are initially used, and longer delays (8s and 12s) are gradually introduced according to a set schedule. This was implemented as we found that during voluntarily performance in the home cage, marmosets would disengage from the operant chamber during longer delay trials if not sufficiently exposed to shorter delay periods. Once animals had completed all delay conditions, working memory maintenance was assessed by fitting an exponential decay function to the mean performance of animals at each delay **(Figure 4I).** Following the method of previous studies (Lind et al., 2015; Nakamura et al., 2018), two parameters were derived from the decay curves to characterize working memory maintenance: the ‘zero-delay performance’, a measure of base working memory performance at an interpolated delay of 0 s, and the ‘performance half-life’, a measure of the rate of working memory decay. In the tested cohort, animals exhibited a mean zero-delay performance of 83.69% (*SD =* 12.05%) and a mean performance half-life of 5.73 s (*SD =* 3.25 s) **(Figure 4J and 4K).** Notably, all animals showed relatively consistent zero-delay performance but a more variable performance half-life, with the working memory of animal M1 decaying at less than half the rate of the other two animals.

### Synchronization of behavioral responses with wirelessly recorded neural activity

Combining home cage-based testing with non-invasive neural recording remains a significant challenge using nonhuman primates, mainly due to the requirement for bespoke technology available in only a handful of research laboratories. Here, we integrated an off-the-shelf system for *in vivo* neural recording with *CalliCog* to demonstrate its broad utility for these applications. A subset of animals that had completed DMTS testing (*n* = 2; M2 and M3) underwent a surgical procedure to implant a radiotelemetry device for ECoG recording. In brief, a microsurgical robot (BrainSight Vet Robot; Rogue Research Inc.) was used to position 4 intracranial electrodes on the surface of the dura mater over the bilateral prefrontal cortex (PFC), using co-registered magnetic resonance imaging/computed tomography (MRI/CT) data to guide placement. The electrodes were insulated with dental cement on the cranium and attached to insulated wires, leading subcutaneously to a radiotelemetry transmitter (HD-S02, Data Sciences International) implanted between the dorsal scapulae. After the surgery, the animals underwent a 14-day recovery period and were reintroduced to home cage behavior testing. As the implant was wholly internalized, neural recordings were non-invasive and acquired via a wireless receiver underneath the operant chamber. The implant was powered on and off before and after behavior experiments using a standard magnet, performed at the cage front without the need for animal handling.

To demonstrate the utility of *CalliCog* for recording synchronous neural and behavioral data, we recorded ECoG from the animals as they performed the DMTS and synchronized this with behavioral timings **(Figure 5A).** First, we used 5 specific timestamps of events in DMTS trials to define 4 trial epochs relevant to working memory processes. This included ‘encoding’, when the animal attended to and interacted with the sample stimulus; ‘maintenance’, the variable delay period when the memory representation was maintained; ‘retrieval’, when the animal retrieved its representation to make a decision; and ‘outcome’, the period in which the animal received either a reward or penalty. Next, as the agent PC that recorded the behavioral data and the telemetry PC that acquired ECoG were synchronized, behavioral timestamps were overlaid over the neural data after the conclusion of experiments. Using custom scripts written in Python, the raw ECoG data went through several preprocessing steps, including bandpass filtering and artifact removal, and was analyzed further offline. We first averaged the processed data from successful trials over each delay condition in the DMTS to confirm that ECoG recordings successfully examined task-related activity. The data was then aligned to the onset of the delay in the maintenance epoch to visualize averaged evoked potentials per animal. Regardless of delay length, voltage changes followed stereotyped patterns, including a smaller peak on or around the delay onset, and a greater, more pronounced peak at approximately 200 ms after the end of the delay **(Figure 5B-5D).** These changes were likely attributed to cortical activity related to the initiation and execution of motor responses, as potentials were evoked regardless of trial outcome (i.e. success or fail) **(Figure 2A-2C).** To determine proof-of-concept for whether ECoG recordings could be implemented to study generalized working memory operations, we performed spectral analysis on the trial data of a single animal M2 and examined ECoG changes across epochs in the DMTS. We selected data from daily recording sessions (*n* = 7) in which the animal performed a minimum of 20 correct trials, and computed the averaged spectra per epoch using a multitaper method **(Figure 5E).** Power was calculated by approximating the area under the resultant power spectral density (PSD) plots across several frequency bands known to be relevant to circuits underlying working memory, including delta (0.5 – 4 Hz, theta (4 - 8 Hz), alpha (8 - 12 Hz), beta (12 - 30 Hz), low gamma (30 - 50 Hz), and high gamma (70 - 100 Hz). The power of each band was then expressed as a percentage of the total power across all analyzed frequencies. Statistical comparisons revealed a distinct re-weighting of relative power between individual epochs that was consistently observed between sessions **(Figure 5F)**. A main effect of the task epoch was detected on the mean relative power across all frequency bands of interest (Friedman test; *p* < 0.01 for all comparisons). In addition, multiple epochs expressed differential power changes within each frequency band. To provide some notable examples of patterns, during working memory maintenance, theta, alpha, and gamma power increased relative to other epochs (Dunn’s test, *p* < 0.05 for all comparisons). During retrieval, delta power increased relative to other epochs, while alpha, beta, and low gamma power were reduced (Dunn’s test, *p* < 0.05 for all comparisons). Alpha power was the most modulated between individual epochs, particularly in a steep reduction between maintenance and retrieval (Dunn’s test, *p* < 0.0001). Finally, the encoding and outcome epochs consistently exhibited the same relative power, regardless of the frequency band of interest (Dunn’s test, *p* > 0.05 for all comparisons). As the purpose of this experiment was to validate the utility of wireless recordings in *CalliCog,* examining the functional causes of these changes is outside the scope of this study. However, these results demonstrate how a third-party neural recording system can be synchronized with *CalliCog* using a simple method to provide definitive and consistent electrophysiological data related to cognitive behavior. The viability of this technique is a promising avenue for future studies to interrogate the neural correlates of cognitive processes using higher powered experimental designs or more targeted recording strategies.

**Figure 5.**
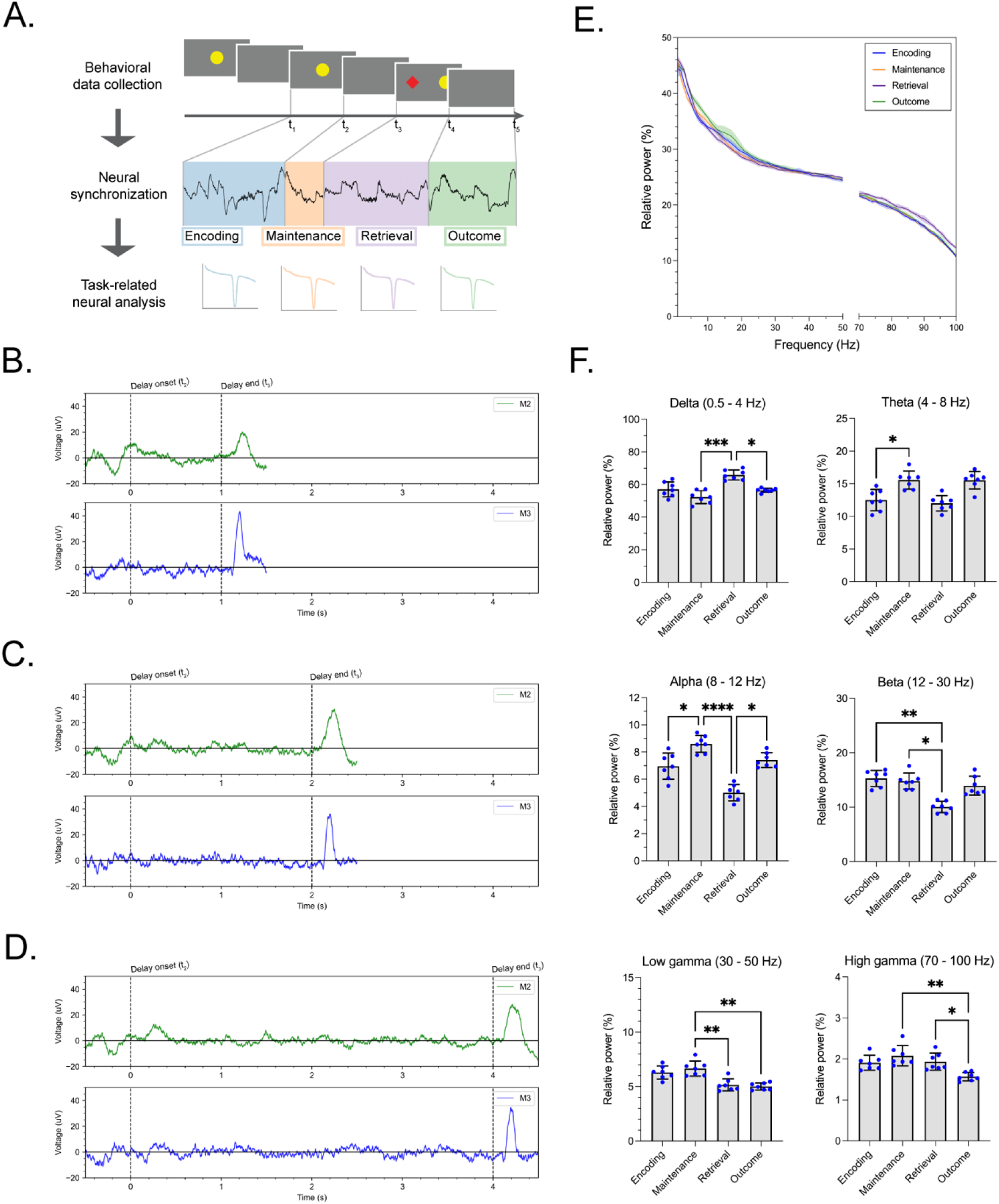
Task-related neural recording in *CalliCog.* **A.** Workflow for task-related ECoG analysis in freely behaving marmosets. Animals (*n* = 2) were implanted with 2-channel ECoG electrodes over the bilateral prefrontal cortex, transmitting live data wirelessly during performance in the DMTS. Five timestamps were recorded from all DMTS trials to define 4 epochs from ECoG data relevant to working memory: encoding, maintenance, retrieval, and outcome. These epochs could then be segmented from the recorded data and analyzed independently. **B-D.** Averaged potentials of filtered, preprocessed ECoG recorded from animals M2 and M3 while performing successful trials at 3 exemplary delay periods of 1, 2, and 4 s (*n* = 70 trials per delay condition). The dotted lines represent the onset and the end of the maintenance epoch. Averaged activity was stereotyped between animals and delay lengths, and evoked events were aligned to the onset of the delay period and following the end of the delay during working memory retrieval. **E.** The averaged power spectra from individual epochs of successful trials from animal M2. Power spectral densities were estimated using a multitaper method, and the data was and log-transformed for visualization purposes. Frequency values between 50 – 70 Hz were included in notch filtering and this excluded from the analysis. **F.** Power quantifications from the power spectra of each epoch at various frequency bands of interest. ECoG power underwent re-weighting between epochs of the DMTS, with all frequency bands differentially recruited in a minimum of one epoch (*p <* 0.05, Friedman test with Dunn’s test for multiple comparisons). Power is expressed as the absolute power of the non-transformed spectrum within each frequency band relative to the absolute power of the entire spectrum. * *p* < 0.05, ** *p* < 0.01, *** *p* < 0.001, **** *p* < 0.0001

## Limitations of Study

A relatively small cohort of animals (*n =* 8) was used in the current work. This number was suitable for the purposes of behavioral testing and validation of the described method, but should not be considered sufficient to account for certain factors in individual variability, including specific age or biological sex. We demonstrated that two subgroups from different housing facilities performed touch training in similar timeframes, indicating that the method was not selective to different housing conditions. However, we caution against the assumption that this will necessarily always apply to marmosets from other colonies or facilities, as behavioral performance could be influenced by several factors including - but not limited to - husbandry environment, diet, social housing, and health status of the colony.

It is important to address that the efficiency of the method used for wireless neural recordings in this study may be influenced by electromagnetic interference (EMI) within an animal facility. Notably, we found that EMI from electronic components in *CalliCog* (e.g. touchscreen and agent PC) was prone to disrupting telemetry frequencies (i.e. between the implant and receiver), and this occasionally caused breaks in the received signal. To mitigate this, we it was imperative to implement shielding and grounding practices to ensure that sources of EMI are contained, not only in the testing device but also in the broader housing area.

Finally, it should be noted that the precision of NTP, used here to synchronize neural recordings and behavioral timestamps, may be variable depending on a user’s configuration. While NTP is capable of sub-millisecond precision, the temporal error of the NTP in the constraints of our setup was sufficient for recording ECoG frequencies but would not be suitable for more transient events recorded by certain techniques (e.g. single-unit recording). Users should consider the reliability of their individual NTP configuration or alternative methods for recording timings if applying this method to different techniques.

## Discussion

*CalliCog* is a groundbreaking system designed to revolutionize cognitive behavioral research with small non-human primates (NHPs) in their home cage environment. This modular, scalable platform allows researchers to conduct automated cognitive experiments effortlessly, harnessing a single interface to control multiple operant chambers within housing facilities. With an intuitive web application and open-source code, *CalliCog* empowers users to create customized behavioral protocols, enabling flexible, hands-off training and testing. Our study with control adult marmosets showcased *CalliCog’s* capabilities, demonstrating how autonomously trained animals excelled in two distinct cognitive tasks. We also achieved a proof-of-concept synchronization of behavioral events with electrophysiological data using wireless ECoG implants, paving the way for new advancements in cognitive neuroscience.

Home cage-based systems for cognitive experiments in NHPs are growing in interest as researchers recognize the advantages for experimental throughput, ethological validity, and animal welfare. While many systems exist, including those made in-house in laboratories or commercially available (e.g., CANTAB cognitive testing systems (Spinelli et al., 2004)), *CalliCog* poses several practical advantages. Firstly, most systems are developed specifically for, or retrofitted from, large NHPs such as macaques and baboons. *CalliCog* was curated for smaller NHP species, such as marmosets, which are experiencing a surge in popularity in cognitive studies due to their suitability for adjunct studies of social cognition (Miller et al., 2016; Samandra et al., 2022) and neurodevelopment (Homman-Ludiye & Bourne, 2020; Homman Ludiye & Bourne, 2016). Resistive touchscreens are conventionally used in large NHP systems due to the force of touch responses. Conversely, we found that a capacitive touchscreen was more likely to register the lighter touch responses of marmosets successfully. Similarly, implementing training bars for the animals to hold for stability was crucial to promoting touchscreen engagement, as unlike larger terrestrial NHPs, marmosets are naturally arboreal, and therefore, less amenable to crouching in flat chambers. In touchscreen tasks, stimulus color was controlled to account for a genetic polymorphism known to affect color discrimination in New World primates (Hunt et al., 1993), which other systems do not consider. A second notable advantage of *CalliCog* is its capacity for automation and ability to adapt to behavioral responses based on single trial responses. While several autonomous systems exist for large NHPs (Berger et al., 2018; Butler & Kennerley, 2019; Cabrera-Moreno et al., 2022; Fizet et al., 2017; Sacchetti et al., 2021; Tulip et al., 2017), only a single instance of a device for marmosets with trial-by-trial functionality has been reported (Calapai et al., 2022). In this study, investigators implemented a robust, dynamic, unsupervised training protocol that adapted to behavioral performance, using sliding windows akin to the rolling window progression criterion implemented in *CalliCog*. However, cognitive tests were notably restricted to discrimination tasks that could be learned through associative learning. Here, we expanded on these findings by demonstrating that tasks implementing abstract rules, as evident by the DMTS, can be likewise achieved by training marmosets using similar protocols. Finally, *CalliCog* is distinct among other systems for its accessibility and flexibility. Non-commercial, freely available systems have been developed for home cage interfaces in NHPs. Still, these have mainly been tailored to the macaque or limited to software applications without the physical infrastructure (e.g. (Hwang et al., 2019)). Here, we provide a complete blueprint comprising the source code, CAD models of hardware, and an inventory of commercially available materials for users to reproduce a working cognitive platform. As demonstrated by our implementation across two independent animal facilities, *CalliCog* is adaptable to different room and cage designs, and the only in-house requirement for users is to mount operant chambers to their cage of choice. Similarly, by modifying the design of the operant chamber, a user could feasibly adapt *CalliCog* for larger NHP species or even integrate the system with testing arenas external to the home cage.

The technical challenge of recording synchronous neural activity while freely behaving NHPs engage in cognitive tasks has been reported previously in macaques (Hansmeyer et al., 2023) and more recently in marmosets (Wong et al., 2023). In both these studies, neural data was recorded or transmitted from devices attached to an external headstage on the animal, which can limit mobility and requires physical restraint to attach. The smaller ECoG telemetry implant described here is wholly internalized and can be remotely activated without imposing the stress of physical handling. This is a considerable benefit to a home cage-based system that leverages behavioral responses without external stressors. However, the recording method described in this study also had notable limitations. We successfully demonstrated proof-of-concept for synchronizing neural recordings with behavioral timestamps in *CalliCog* and demonstrated that averaged ECoG activity was aligned with task events in the DMTS. In addition, 2-channel ECoG recordings were capable of capturing reproducible activity relevant to functional epochs within the working memory task, though a direct link between these changes and generalized working memory processes warrants further investigation. As the recording method used here was a novel approach for studying working memory in NHPs, conclusions regarding how these findings compare to existing data are limited. To date, most studies examining working memory activity in NHPs used intracortical electrodes or arrays to record local field potentials (LFPs) in the PFC, mainly in macaques. Summed activity from these LFP recordings showed task-related changes in amplitude comparable to the averaged potentials investigated in this study, with peaks in activity aligned to both the delay onset and during trial reward (Holmes et al., 2018). Therefore, PFC activity from ECoG was reliably recruited in a task-dependent manner to what has been previously described. However, our findings from the spectral analysis diverged from that canonically observed through LFP in working memory. Previous work demonstrated a recruitment of alpha/gamma and a reduction in beta power during memory encoding (Holmes et al., 2018; Lundqvist et al., 2016; Wang et al., 2022), and transient yet recurrent beta/gamma activity during maintenance (Lundqvist et al., 2016; Miller et al., 2018). Our analysis of spectral power between working memory epochs did not appear to support this distinction, for at least two likely reasons: temporal changes in power were not considered in our analysis, thus losing the resolution to observe transient events; and the use of surface electrodes lacks the precision to record from working memory-modulated sites. Indeed, the results from spectral analysis in this study more closely align with that from clinical electroencephalography (EEG) studies. For example, among the most reproducible findings from clinical EEG is the elevation of prefrontal gamma and theta activity during maintenance, which are thought to contribute differentially to information stored in working memory (Roux & Uhlhaas, 2014). The evidence in our study that gamma and theta power were also reliably enhanced during memory maintenance supports this functional association. Despite this, the purpose of this study was to demonstrate feasibility of the technique, and more studies with higher throughput are required to make further conclusions. Instead, here we demonstrate the power of synchronized neural recordings for examining the physiological correlates of behavior. ECoG signatures of individual epochs within single trials were notably reproducible across sessions despite the natural variability of behavior in an unrestrained environment. This approach, therefore, constitutes a useful tool for examining dynamic states of cognitive behavior, which could be leveraged in future studies to explore mechanisms relevant to cognitive disorders. Similarly, the process for synchronization is not restricted to ECoG, and could be applied to more precise recording techniques such as multi-channel single-unit electrophysiology or intravital miniscopes for optical imaging.

Both reversal learning and working memory have been extensively studied in marmosets, including in home cage settings. Here, we selected experimental designs adapted from previous studies (Nakamura et al., 2018; Takemoto et al., 2015) to benchmark the performance of control marmosets undergoing automated training against data from conventional methods. In the reversal learning task, marmosets performed an average of 85.8 errors to criterion, compared to approximately 50 – 100 errors on average in the literature for young adult animals following the same proficiency criterion (Munger et al., 2017). This suggests that the performance of animals using *CalliCog* could reliably replicate that attained using manual methods. In the DMTS, individual marmosets in this study demonstrated the capacity to perform above chance level for at least the longest delay length of 12 s, comparable to those in the literature. However, on average the cohort exhibited relatively worse performance as the delay length was increased (Nakamura et al., 2018). While this could be attributed to the testing environment, the version of the DMTS implemented here notably selected from 3 possible stimuli during sample presentation rather than the 2 commonly used in delayed response tasks. This difference, while encouraging marmosets to respond more frequently to task rules than recency bias, likely increased the difficulty and, therefore, impacted overall performance.

In conclusion, using an automated system, we present a method for home cage-based cognitive behavioral experiments in small NHPs. Behaviorally naïve marmosets were trained to proficiency in separate cognitive tasks using an automated protocol, and task-related wireless ECoG was recorded during working memory tasks. By providing this openly available method, we endeavor for cognitive testing in NHPs more accessible for members of the scientific community who are otherwise hindered by lack of resources or expertise with NHP behavior. This has the potential to encourage more studies of primate cognition, improving translational insights into primate-specific circuits and behaviors implicated in both health and disease in humans.

## Resource availability

### Lead contact

Requests for further information and resources should be directed to and will be fulfilled by the lead contact, James Bourne.

### Materials availability

This study did not generate new unique reagents.

### Data and code availability

- Behavioral and electrophysiological data required for the analyses have been deposited at Figshare and are publicly available as of the date of publication via the DOI listed in the key resources table.
- A snapshot of the original code for running the *CalliCog* system and all behavioral experiments in this publication has been deposited at Figshare at the time of publication at the link provided in the key resources table. The lead author can provide custom code for the analysis of electrophysiological data upon request.
- Any additional information required to reproduce the data reported in this study is available from the lead contact upon request.

## Acknowledgements

The authors thank Felix Sng, Kieren Pinto, and the NIH Section on Instrumentation for engineering assistance; and Leor Katz, Kevin Marche, and Terry Chow for providing input on the analyses. We also extend our thanks to Monika Toth, Annabel Estasy, and Xianfeng (Lisa) Zhang for providing care and support for research animals. This research was supported by the National Health and Medical Research Council, a NARSAD (Brain & Behaviour Research Foundation, US) Independent Investigator Award, and the Intramural Research Program of the NIMH (ZIA MH002984) to J.A.B. The Australian Regenerative Medicine Institute is supported by grants from the State Government of Victoria and the Australian Government.

## Author contributions

J.T.S. performed data collection and analysis; J.T.S., B.L.M.V., B.J.S. and D.K.J.R. developed the software; J.T.S., B.L.M.V., D.D.P., and A.Y.F. designed and built the hardware; J.A.B. provided general supervision; J.T.S and J.A.B. conceived the project and wrote the paper.

## STAR Methods

### Key resources table

3 Key resources table
| REAGENT or RESOURCE | SOURCE | IDENTIFIER |
| --- | --- | --- |
| **Experimental models: Organisms/strains** |  |  |
| Common marmoset (*Callithrix jacchus*) | National Nonhuman Primate Breeding and Research Facility, Australia.<br>National Institute of Mental Health (NIMH) | N/A |
| **Software and algorithms** |  |  |
| *CalliCog* source code | This paper | Snapshot from date of publication (v1.0.0):<br>https://doi.org/10.6084/m9.figshare.27873153<br>Actively maintained repository:<br>https://github.com/NIMH-SCCN/callicog |
| Python (v3.8.19) | Python Software Foundation | www.python.org |
| PsychoPy (v2021.2.3) | Open Science Tools Ltd. | www.psychopy.org |
| PostgreSQL (v15) | The PostgreSQL Global Development Group | www.postgresql.org |
| RPi-Cam-Web-Interface (v6.6.15) | silvanmelchior | https://elinux.org/RPi-Cam-Web-Interface |
| Ponemah (v6.51) | Data Sciences International | www.datasci.com/products/software/ponemah |
| MNE-Python (v1.7.0) | MNE Developers | https://mne.tools/stable/index.html |
| MATLAB | The MathWorks, Inc. | www.mathworks.com |
| cp_wrapper() | Gallistel et al. (2004) | https://doi.org/10.1073/pnas.0404965101 |

| **Other** |  |  |
| --- | --- | --- |
| *CalliCog* inventory | Various sources | https://doi.org/10.6084/m9.figshare.27873153 |
| PhysioTel HD-S02 radiotelemetry implant | Data Sciences International | 270-0173-001 |
| Matrix 2.0 (MX2) | Data Sciences International | 271-0119-002 |
| RSC-1 radiotelemetry receiver | Data Sciences International | 272-6012-001 |

#### Experimental model and study participant details

A total of 8 adult marmosets (*n* = 8; 3 female, 5 male) aged a minimum of 12 months were used in the current study. Biological sex was not a selection criterion chosen for the cohort, and the experimental cohort was randomly selected based on animal availability. All animals were naïve to behavioral experiments, but all except 2 (M7 & M8) had previously undergone MRI scanning under anesthesia for an unrelated study (maximum of 2 scans per animal; scan time < 90 min). Within the current study, some animals were excluded from certain experiments due to transferal to or from other adjunct projects. For complete details of sample sizes per experiment, refer to **Table S1.**

Housing of the experimental cohort was split between 2 individual facilities assigned as Facility 1 (M2, M3, M7, and M8) and Facility 2 (M1, M4, M5, and M6). Facility 1 was an indoor vivarium and housed vertically stacked double cages joined by a tunnel (200 cm x 78 cm x 78 cm, 13 cm elevation). Facility 2 housed singular cages (144 cm x 75 x 64 cm, 40 cm elevation) equipped with controllable access tunnels to a larger outside enclosure, which animals could freely explore during 09:00 – 14:00 when not under experimentation. All animals were reared by their parental dyad and were not relocated from their family unit to separate housing until at least 12 months of age. Most animals were single-housed, but retained visual and audial access to other marmosets in the vivarium. The exception to this is animal M8, which was pair-housed with a sibling and was restricted to one half of the cage during in-cage testing. Both Facility 1 and 2 operated on a 12-hr light/dark cycle (07:00 - 19:00) and controlled ambient conditions (26 - 28 °C, 40 – 60 % humidity). Animals were fed a rotating diet comprising various fruits, nuts, insects and formulated primate pellets or chow once daily after experimentation, and had *ad libitum* access to water at all times. In-cage behavioral testing was performed between 09:00 – 17:00 daily from Monday to Friday, with each individual animal provided access to an operant chamber for between 3 to 6 hours per day, depending on schedule and engagement.

All animal procedures in Facility 1 were fully compliant with the Guidelines for the Care and Use of Laboratory Animals by the National Institutes of Health and approved by the Animal Care and Use Committee of the National Institute of Mental Health. All animal procedures in Facility 2 were conducted in accordance with the Australian Code of Practice for the Care and Use of Animals for Scientific Purposes and were approved by the Monash University Animal Ethics Committee, which also monitored the welfare of the animals.

### Method details

#### Touchscreen training

Animals were trained to operate the operant chamber touchscreen by progressing through 7 task phases designed to optimize accuracy and mitigate response bias. In phase 1, trials consisted of a single full-screen stimulus pseudorandomly sampled to be red, yellow, or blue (RGB) in color. This mitigated color bias, as red-green dichromats (all male marmosets) can equally discriminate them (Hunt et al., 1993). Touching the stimulus on a given trial resulted in positive reinforcement (∼150 ul liquid reward, 800 Hz pure tone). In phase 2, trials consisted of a square stimulus (700 x 700 px) presented centrally. Here, animals learned to associate positive and negative outcomes by targeting the stimulus and avoiding the grey background, which was associated with a penalty (2s timeout period, 100 Hz tone). During phases 3 – 5, response accuracy was shaped by gradually reducing the stimulus size in 150 x 150 px steps until the target size of 250 x 250 px was reached in phase 5. Finally, during phases 6 and 7, stimuli were presented randomly on each trial, firstly within a smaller area (540 x 260 px) and then at any position on the screen. This encouraged animals to respond to stimuli equally regardless of spatial position. For phases 2 – 7 of touchscreen training, animals progressed to the subsequent phase after achieving a session-based progression criterion of 80% success in 3 consecutive sessions of 50 trials. For phase 1, animals progressed either at this criterion or after an investigator had determined that stimulus-reward association had been reliably established.

#### Novel discrimination and reversal learning

Before testing in novel discrimination and reversal learning, animals underwent discrimination training. During training, animals respond to a cue stimulus (black square) to initiate a trial. Following a touch response, 2 novel stimuli were pseudorandomly presented in left and right positions. Stimuli consisted of geometric images composed of the 3 test colors on a white square background and were deterministically associated with either reward or penalty outcomes according to a set schedule. Animals were granted a period of 2 s to respond to either stimulus before the trial ended. Touches recorded outside of the stimuli were inconsequential and resulted in neither reward nor penalty. Animals performed 3 iterations of discrimination training and learned to discriminate the correct from the incorrect stimulus in each pair. Progression to each subsequent phase occurred after achieving a session-based progression criterion of 80% success in 3 consecutive sessions of 50 trials. Upon completion of discrimination training, animals performed a further 10 novel discrimination experiments with an identical structure to the training phases, each incorporating a novel stimulus pair. Each discrimination experiment was associated with a reversal learning phase, initiated after animals had reached a session-based progression of 80% success in 3 consecutive sessions of 50 trials. During reversal learning, the stimulus-reward contingencies of each stimulus were reversed, and animals performed the task to a rolling average progression criterion of 90% success over a 100-trial rolling window.

#### Delayed Matech-to-Sample (DMTS)

Testing in the DMTS was preceded by 4 individual training phases designed to approximate the desired behavior in the final task successively. During phase 1, trials began by presenting a ‘sample’ stimulus in a central position, pseudorandomly selected to be either a yellow circle or a red diamond. As the stimuli differed in shape and area, they were overlaid with invisible ‘hit boxes’ of an equal 250 x 250 px to ensure that animal responses to the stimuli were not affected by variability in precision. Responding to the sample stimulus initiated a 0.5 s delay followed by a forced choice window including the sample and a distractor (the remaining stimulus not selected as the sample) in left and right positions. Animals were granted a period of 3 s to respond to either stimulus before the trial ended and received reward or a timeout penalty for selecting the sample or distractor, respectively. In phase 2 of DMTS training, trials were mainly identical to phase 1, but in addition the sample was presented twice at the start of the trial to ensure animals visually attended to it. In phase 3, trials were mainly identical to phase 2 but now included a center position as a possible location for a stimulus to appear during forced choice. In phase 4, a third stimulus, a blue star, was added to the pool of stimuli that could be selected as either the sample or distractor on a given trial. We intentionally used 3 stimuli in the pool rather than 2 to optimize experimental validity, as nonhuman primates are known to be heavily biased in behavioral tasks by stimuli that were rewarding in recent trials (i.e., the recency effect (Brunet & Jagadeesh, 2019). Throughout all DMTS training, animals progressed to each subsequent phase after achieving a session-based progression criterion of 80% success in 3 consecutive sessions of 50 trials.

Once trained, animals performed DMTS testing. Trials were identical to the final training phase, except that more extended delay periods were now pseudorandomly introduced between the final presentation of the sample stimulus and forced choice. As we determined during pilot studies that longer delays would result in animals leaving the operant chamber and terminating the trial early, engagement was sustained by successively introducing longer delays as the animals completed greater numbers of trials. Using target-based progression criteria, animals initially performed trials including delay conditions of 0.5, 1, 2 and 4 s, and this was incrementally extended to include an 8 s and then 12 s condition. In total, animals performed 400 trials of 0.5 – 4 s delays, 300 trials of the 8 s delay, and 150 trials of the 12 s delay conditions.

#### Behavioral data analysis

For the reversal learning task, averaged learning curves were generated per animal by calculating the trial-wise average from binary performance outcomes (i.e. success or fail). The resultant probability values were smoothed for visualization purposes over a rolling window of 100 neighbors using a 2^nd^ order polynomial as per a previous method (Savitsky & Golay, 1964). Next, we applied a different method for learning curve analysis (Gallistel et al., 2004) previously used in several marmoset studies (Sadoun et al., 2019; Takaji et al., 2016; Takemoto et al., 2015) to group trials into discrete stages of learning. The binary performance data from each reversal learning experiment were input into a recursive algorithm that detected ‘change points’ in the frequency of cumulative successes using χ^2^ tests. A trial was designated as a change point if the frequency of cumulative successes at that point was greater than expected by chance, as determined by a criterion of *p* < 0.02 (logit = 1.7; for more detail, see previous studies (Sadoun et al., 2019; Takemoto et al., 2015)). The change points were then used to segregate the learning curve into 3 stages of reversal learning: ‘perseverative’, ‘learning’, and ‘achieving’. Performance during these stages was calculated as the averaged cumulative performance across trials within each stage. In addition, we calculated the likelihood of an animal exhibiting win-stay or lose-shift strategies during the learning phase of each reversal learning experiment, which reflects the ability to learn the new stimulus-reward contingencies based on positive or negative feedback. The probability of win-stay trials was defined as the ratio of successful trials to failed trials immediately following a successful trial, and the probability of lose-shift trials as the ratio of successful trials to failed trials immediately after a failed trial. These probabilities were averaged over all reversal experiments for comparison between animals.

For the DMTS, the primary metric for working memory function was the average success rate per delay condition. To examine the relationship between working memory performance and delay condition, we used a method for working memory analysis (Lind et al., 2015) applied in a previous marmoset study (Nakamura et al., 2018). The performance data over delay length for each animal was fit with a simple exponential decay function:

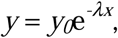

where *λ* is the decay constant and *y_0_* represents the zero-delay performance, or the estimated success rate at a delay of 0 s. The estimated zero-delay performance was used to quantify the baseline ability of an animal to perform the working memory task competently (i.e. ability to match two identical stimuli). The performance half-life was also calculated from the decay function by examining the delay time at half the duration between maximal performance and chance (50% success). It was used to quantify the capacity to maintain working memory over time. For visualization purposes, the decay function presented was the mean of the individual decay curves calculated for each animal, with the upper and lower boundaries representing standard deviation. The plot also depicts a threshold for performance not statistically different from chance level, as calculated via a 2-tailed binomial test based on the sample size of 0.5 – 4 s delays (*n* = 400 trials).

#### ECoG implantation surgery

Anesthesia was induced by an intramuscular injection of alfaxalone (8 mg/kg; Jurox) combined with the sedative diazepam (3 mg/kg; Pfizer) and maintained with inhalant isoflurane (1 – 2% in oxygen). Atropine (20 µg/kg; Pfizer) and dexamethasone (0.2 mg/kg; Ilium) were also administered intramuscularly to reduce mucosal secretion and minimize cerebral edema, respectively. From the moment of induction, anesthesia level, heart rate, blood oxygen saturation, and external temperature were constantly monitored and kept within optimal ranges throughout the procedure. Once anesthetized, the scalp and upper back were shaved, and the animal was secured to a stereotaxic frame. Local anesthetic (0.3 ml bupivacaine; Aspen) was delivered subcutaneously to the surgical sites, and the sites were disinfected three times with 10% iodine solution (Betadine). Under aseptic conditions, a horizontal incision was created between the scapulae, and a subcutaneous pocket was formed posterior to the incision. A radiotelemetry transmitter (HD-S02; Data Sciences International; 28 x 17 x 8 mm) was inserted into the subcutaneous pocket, and wires were routed subcutaneously to a second surgical incision at the posterior aspect of the scalp. The dorsal incision was then sutured (4-0 Vicryl Rapide, Johnson & Johnson), and the scalp incision lengthened along the midline to the anterior aspect of the scalp. With the cranium exposed, the skull surface was registered with an optical laser driven by a microsurgical robot (BrainSight Vet Robot; Rogue Research Inc.) to structural MRI/CT data, which was acquired from the surgical animal and co-registered less than 1 week prior. Using the structural MRI and a digital stereotaxic atlas of the marmoset brain (Majka et al., 2020; Paxinos et al., 2012) as a reference, 2 intracranial electrodes were placed in each hemisphere along the anteroposterior axis above the PFC. Electrode placement was aligned to stereotaxic coordinates (16.5 mm and 15.5 mm anterior to the interaural line, +/- 2 mm lateral to the midline) and then adjusted slightly for brain morphology so that the electrodes spanned areas 9, 46D, 46V, 8b, and 8aD. For electrode placement,1.4 mm burr hole craniotomies were performed at each position, and iridium oxide screws were advanced into the bone with the ends contacting the dura mater. Wires from the telemetry device were then anchored to the screws, and the screws were insulated with dental cement (C&B Metabond, Parkell Inc.). The scalp incision was then sutured, and the animal was recovered from anesthesia over a homeothermic blanket with 100% medical oxygen. Post-operative analgesic was administered at 72 hrs post-surgery.

#### ECoG recording, post-processing, and analysis

ECoG was recorded from implanted animals performing the DMTS via a telemetry receiver placed underneath the operant chamber. 2-channel ECoG (1 channel per left and right PFC) was acquired at a sampling rate of 500 Hz and visualized in Ponemah software (version 6.51; Data Sciences International). The raw ECoG data was then exported and processed offline using custom Python scripts and functions from the package MNE-Python (version 1.7.0) (Gramfort, 2013). Firstly, behavioral data was exported manually from *CalliCog,* and timestamps were aligned with the raw ECoG recording. Next, data breaks from signal dropout were detected when animals moved away from the telemetry receiver (e.g. while entering and exiting the chamber) and these were excluded from further analysis. Similarly, we performed thresholding on spectrograms of raw data to identify substantial, transient increases in broadband power resembling contamination from muscle contractions, which typically occurred while animals were feeding from the reward spout. Trials containing data with this artifact were excluded from subsequent analyses. The valid data was then fed through a notch filter at 60 (+/- 5) Hz for line noise removal, and bandpass filtered from 0.5 to 100 Hz with a zero-phase, sixteenth-order Butterworth filter. The data from both channels was then averaged for all further analyses.

To visualize evoked changes in ECoG activity across trials, the averaged potentials of processed data were generated by querying the datasets for all trials meeting certain conditions (i.e. delay length, successful or failed outcome) and then aligning data from each trial to the onset of the maintenance epoch delay. The trials were then averaged and depicted as plots of the mean +/- standard error. We performed spectral analysis on the individual epochs of all valid, successful trials to examine changes in ECoG spectral power. Firstly, trial data was segmented into four epochs that correlated with behavioral timestamps from each trial, as follows:

- ‘Encoding’: Released touch from the first sample window to the released touch of the second sample window.
- ‘Maintenance’: Released touch of the second sample window to the presentation of stimuli in the forced choice window.
- ‘Retrieval’: Presentation of stimuli in the forced choice window to the touch of a stimulus.
- ‘Outcome’: Touch of a stimulus in the forced choice window to the beginning of the subsequent trial.

Next, power spectral densities were estimated from individual epochs of trials using a multitaper method with 4 Slepian tapers, 624 ms windows, and a time half-bandwidth product of 2.5. These power spectral densities were then averaged across individual recording sessions (*n >* 20 trials per session), and transformed for visualization purposes using the following equation:

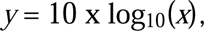

where *x* is the original power spectral density (µV^2^/Hz) and *y* is the log-transformed (dB). Next, power was calculated, per epoch, by estimating the area under the curve of the non-transformed mean power spectral densities between the ranges of specific frequency bands of interest: delta (0.5 - 4 Hz), theta (4 - 8 Hz), alpha (8 - 12 Hz), beta (12 - 30 Hz), low gamma (30 - 50 Hz), and high gamma (70 - 100 Hz). This was achieved using Simpson’s rule for approximating integrals using parabola, following a method from the Python package YASA (Vallat & Walker, 2021). Band power calculations were then normalized to the computed power of the entire spectrum for statistical comparisons.

### Quantification and statistical analysis

All statistical tests were performed in Prism 10 (GraphPad) using a significance criterion of 0.05. As most data violated the assumption of normality due to small sample sizes (D’Agostino-Pearson test; *p* < 0.05), only nonparametric statistical tests were used. A Mann-Whitney U test was used for all pairwise comparisons, and a Kruskal-Wallis test or Friedman test was used with Dunn’s test for multiple comparisons for all group-wise comparisons. In the analysis of touchscreen and discrimination training, Z-score calculations were used to identify outlier data, with the range outside -2 to 2 treated as the threshold for outliers. For all behavioral results, the sample size (*n*) was defined as the performance per animal or mean performance per animal, or for comparisons between animals (e.g. errors to criterion) defined as individual session performance per animal. Visual representation of variation is represented by standard deviation in bar charts, and interquartile ranges in box plots.

---

**Figure S1.**
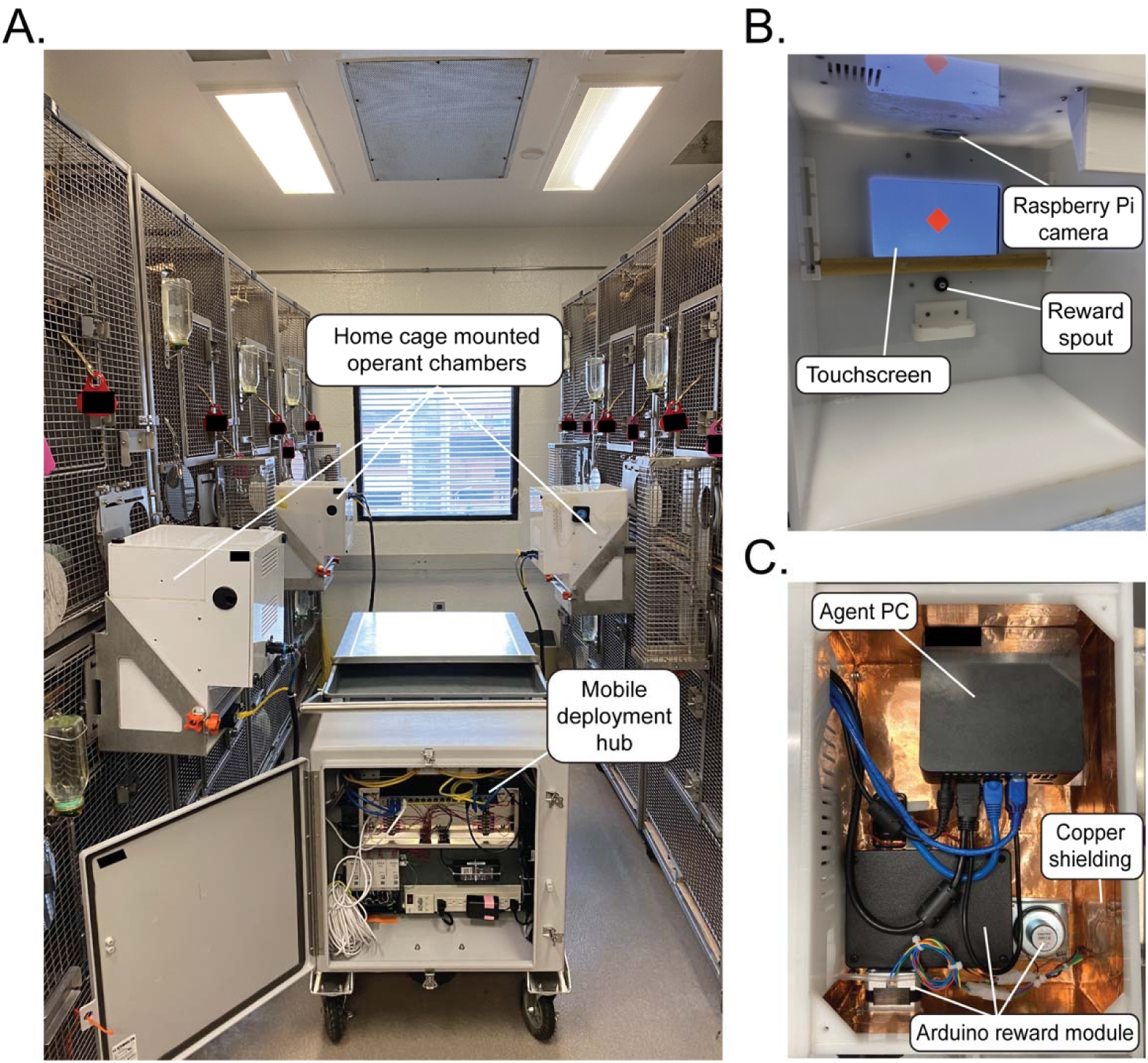
Deployment and composition of operant chambers in *CalliCog.* **A.** Operant chambers are mounted to individual home cages in an animal facility using custom-made brackets. Each individual unit is supplied with mains power and an ethernet connection to a local network. In the configuration pictured, this was facilitated by a custom-made mobile deployment hub (not required to run *CalliCog*) containing a network switch, router and a telemetry interface, which was also used to wheel operant chambers into the housing area for deployment. Note: Home cages in this facility comprised two enclosures stacked vertically and joined by external tunnels, which were used to temporarily separate pair-housed animals for cognitive testing. **B.** Internal view of the operant chamber. The animal interface comprised a capacitive touchscreen, reward spout and tray, and a wooden training bar that could be adjusted in height for animals to hold during task performance. Animals were visually monitored via a Raspberry Pi camera module embedded in the chamber ceiling. **C.** The upper compartment of the operant chamber houses the agent PC, Arduino reward module, and Raspberry Pi camera module. To limit electromagnetic interference interrupting wireless telemetry recordings, copper shielding lined the surface between the devices and the internal chamber, and both devices and shielding shared a common ground.

**Table S1.** Allocation of experimental animals. A subset of animals (M4 – M8) were transferred to adjunct projects after novel discrimination and reversal learning, and did not participate in the DMTS. Only 2 animals (M2 and M3) were selected for ECoG implantation and recording. **\***Animal M5 underwent touchscreen training using a slightly modified protocol, so data from this animal was not included in the results comparing performance between animals. ** Animal M3 only completed 3 novel discrimination and reversal learning experiments before being fast-tracked to DMTS training for another study, so data from this animal did not contribute to the results for novel discrimination or reversal learning.

| **Animal** | **Facility** | **Sex** | **Age at training onset** | **Touchscreen training** | **Novel discrimination and reversal learning** | **DMTS** | **ECoG** |
| --- | --- | --- | --- | --- | --- | --- | --- |
| **M1** | 2 | F | 1 y, 5 mo. | X | X | X |  |
| **M2** | 1 | M | 4 y, 11 mo. | X | X | X | X |
| **M3** | 1 | M | 2 y, 7 mo. | X | ** | X | X |
| **M4** | 2 | M | 2 y, 8 mo. | X | X |  |  |
| **M5** | 2 | F | 1y, 0 mo. | * | X |  |  |
| **M6** | 2 | M | 1 y, 0 mo. | X | X |  |  |
| **M7** | 1 | M | 2 y, 8 mo. | X | X |  |  |
| **M8** | 1 | F | 1y, 0 mo. | X | X |  |  |

**Figure S2.**
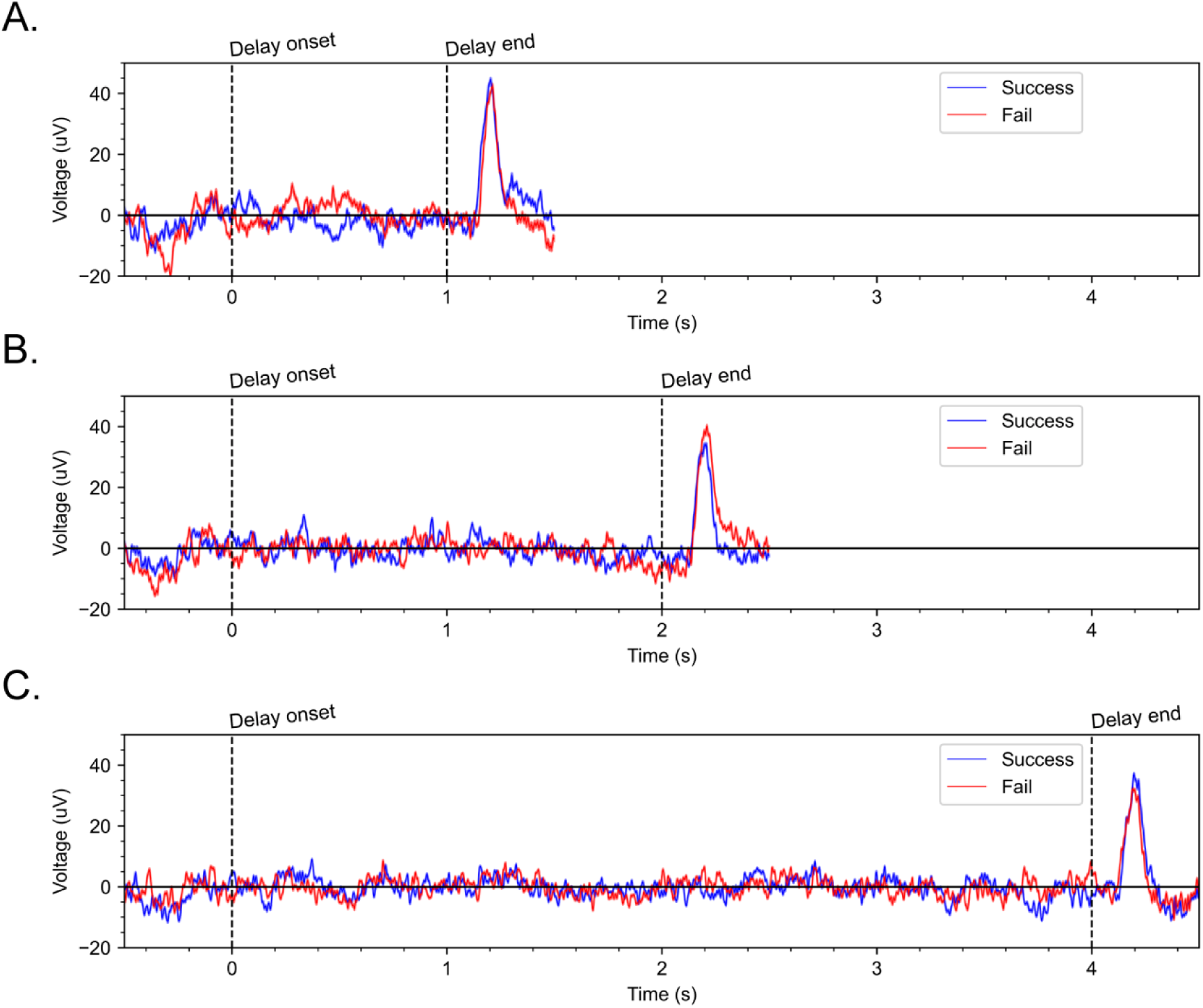
Examples of averaged potentials from the preprocessed ECoG of animal M3 during the delayed match-to-sample task. **A-C.** Averaged potentials from either successful or failed trials across 3 different exemplary delay periods of 1, 2, and 4 s (*n* = 40 trials per condition). Activity appeared to be relatively stereotyped despite differences in trial outcome, particularly at approximately 200 ms following the end of the delay (second dashed line).

